# Diffractive scanning live volumetric two-photon microscopy within the contracting mouse intestine

**DOI:** 10.64898/2026.03.18.712419

**Authors:** Jonas Jurkevičius, Milvia Alata, Moritz Wiggert, Maximilian Rixius, Stefan Reinhards, Matthea Thielking, Christian Stock, Candice Fung, Alexandre Favre, Dirk Theissen-Kunde, Luigi Bonacina, Sebastian Karpf, Pieter Vanden Berghe

## Abstract

Obtaining structural information from the enteric nervous system (ENS) within intact intestinal tissue requires microscopy systems capable of imaging through multiple tissue layers and during ongoing physiological motion. Tissue opacity, three-dimensional geometry, and spontaneous contractions strongly constrain volumetric imaging, limiting the applicability of most conventional linear optical techniques to imaging in either dissected, stretched or pharmacologically suppressed tissues. We apply Spectro-temporal Laser Imaging by Diffracted Excitation (SLIDE) microscopy, a diffraction-based scanning approach enabling fast volumetric two-photon imaging, to record the ENS in an intact *ex vivo* intestinal preparation from a transgenic mouse line expressing the red fluorescent protein TdTomato in peripheral and enteric neurons and glia. We achieved fast volumetric imaging during spontaneous contractions, capable of resolving micrometer-scale displacements in three dimensions, without inducing observable photodamage or compromising tissue viability over the experimental timescale. This work establishes 4D-SLIDE microscopy as a robust experimental framework for visualizing enteric neural structures within their native three-dimensional context during physiological motion, with direct relevance for conditions involving altered intestinal mechanics.

## I. INTRODUCTION

The ability to extract spatial and temporal information at the relevant length and time scales is essential for understanding biological processes. To date, no single microscopy technique can address all the challenges associated with physiological imaging in complex tissues. Although robust solutions exist for individual constraints, none of the current approaches resolve them simultaneously. Widefield systems enable image acquisition at kilohertz rates but lack intrinsic three-dimensional sectioning^1^; light-sheet microscopy is restricted to optically transparent specimens^2^; and two-photon microscopy, while offering substantial penetration depth, becomes inherently slow when volumetric imaging with adequate spatial sampling is required^3^. Here we take advantage of the diffractive 2-photon scanning technique developed by Karpf et al.^4^ and demonstrate the operation of a biology lab grade microscopy setup to capture volumetric data in living contractile tissues. As a key example, we image the enteric nervous system (ENS) embedded in between the contractile muscle layers of the mouse small intestine.

The gastrointestinal tract is a crucial organ that provides nutrients and molecular building blocks to the organism. It is shaped by evolution into a tubular organ, with different compartments that allow passage (the intestines) and temporal storage of food (the stomach) and waste material (the large intestine). The two muscle layers (one circumferential and one longitudinal) in the intestinal wall serve to grind, mix and move the food contents through the entire tract. This so-called gut motility results from coordinated actions of the enteric nervous system (ENS), a dual layered neuro-glia network that is embedded in the intestinal wall and that coordinates muscle contractility, bloodflow, and secretion. The latter two functions are controlled by the submucous plexus (SMP), while the muscle and motor control is tasked to the myenteric plexus (MP), embedded in between the two muscle layers^5^. Circular muscle contraction narrows the digestive tube pushing food forward towards locally relaxed areas, while the longitudinal muscle contraction shortens the intestine at the same time. Together, this generates a peristaltic propulsion of the food contents. In the small intestine, the movement of these muscle layers enables digestion and, by secretion of water, also absorption of nutrients. At the time most nutrients have been depleted from the lumen, the contractions mainly serve to push undigested materials further along the intestine to finally be removed from the organism.

Overall, intestinal motility has been studied using various approaches, including manometry, spatio temporal mapping, fluoroscopy and MRI and *in vivo* quantification of electrolyte and fluid flux^6–8^. These approaches mainly provide data on the overall function or describe the total composed movement. However, in order to understand how the ENS controls all these functions, live microscopy is required. Recently, we were able to show that different neuronal populations respond differently to specific nutrients as they were exposed to the tip of the intestinal villi^9^. Importantly, communication between the different layers, epithelial, mucosa, submucosal, SMP and MP neurons is crucial to enable proper functioning^10^(Fig.1 b).

**FIG. 1:**
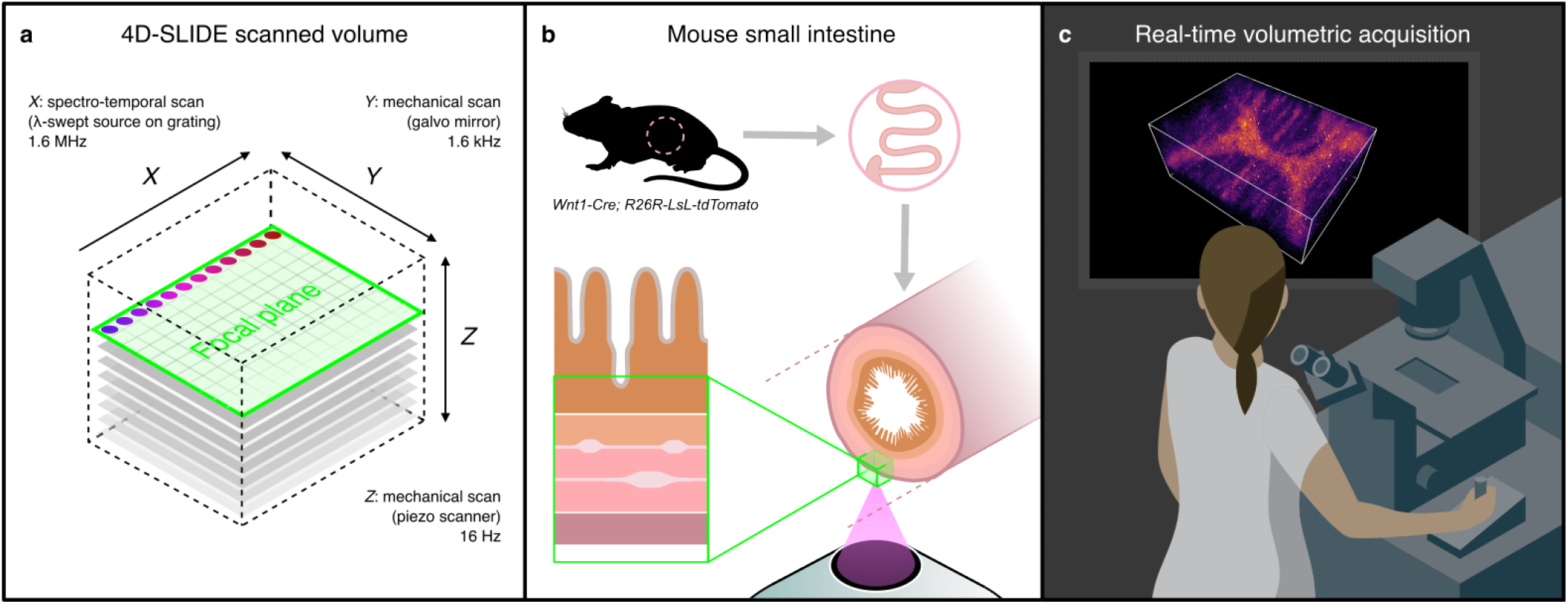
Schematic overview of the 4D-SLIDE system used in this study to image inside the living contracting intestine. **a** Three-dimensional raster scanning for 16Hz volume-rate acquisitions (512x1024x100 voxels). **b** The small intestine was excised from the mouse and the enteric nervous system of living intestine was recorded on an inverted microscope. The acquired volume (in green) contains multiple neuron layers and the mucosal crypts. **c** During the experiment, a real-time 3D live render of the 6.8GByte/s data stream is displayed on the computer screen, enabling fast interactions and 3D navigation within the living sample.

Even though widefield imaging can extract important physiological information about ENS function and similarly, confocal and classic 2-Photon imaging have been used to derive structural composition of these different layers, no technique exists that captures the fast moving components in the gut wall with sufficient resolution and at sufficient speed to further our understanding of the connections between the SMP and MP layers. To this date, no one has been able to investigate the effects of gut (longitudinal and circular) contractions on the two layers of nerves separately, because imaging systems were either too slow or were not able to optically section and penetrate deep enough into the tissue to generate a volumetric dataset with adequate informational content. Especially when tissues are alive, and thus contractile, no microscopy technique is able to track the 3D movement of MP and the SMP in real time. As the forces and stretches on these neuronal processes must be enormous, at least from a cellular perspective, they may well be relevant with respect to physiological control of function, or be a source of discomfort in case of unphysiological contractions.

The ability to record these movements (minute at the organ- and major at the cell scale) is key to unravel the details of ENS and gut function. In this work, we present the first physiology application of 4D-SLIDE microscopy and demonstrate its ability to image inside living contractile tissue and detect micrometer-scale distortions in 3D. We show that this novel imaging modality can be used without causing observable damage or a detrimental impact on its physiology.

## II. MAIN

### 4D-SLIDE Imaging of Living Tissue

For high-speed volumetric two-photon microscopy of living mouse intestine, we designed a dedicated 4D-SLIDE microscope built as a rack unit containing all components, making the system moveable and robust. The scanning details, schematic depiction of volumetric scanning in a whole intestinal tube, and real time visualization by 4D-SLIDE system are presented in Figure 1. Our goal was to simultaneously image both ENS layers in a contracting intestinal tube (Fig. 1 b). To achieve the required fast volumetric acquisition speed, (Fig. 1 c) we used a swept-source FDML-MOPA laser^11^ to deliver pulses of 30 ps duration at 802 MHz repetition rate, corresponding to 512 pulses per sweep or 512 imaging pixels along the *X*-axis. In our experiment, the line-scan rate was set to 1.6 MHz (Fig. 1a), equating the laser sweep rate. The galvo mirror for y-axis scanning was operated bi-directionally for 1.6 kHz frame-rate (1024 lines per image). The axial dimension was scanned using a piezo objective scanner for 16 Hz volume rate (100 imaging planes per volume). The 16 Hz rate volume acquisitions were processed by custom software and rendered in real-time in *Napari*^12^. Due to the high repetition rate and the picosecond excitation pulses, SLIDE employs higher average power than usual in femtosecond 2-photon systems. Consequently, we first checked the compatibility of the Watt-level excitation for living tissues. These experiments, in freshly dissected intestinal tissue (stretched gut tissue preparation, see methods) immobilized over an INOX ring, were performed to examine bleaching and phototoxicity at varying excitation powers. Four different regions were imaged for ∼ 60 s, each with different excitation powers (see Table I) . The lowest power used, 100 mW (region I), generated no noticeable bleaching or visible photodamage but also poor image quality (Fig. 2a). In region IV, exposed to the highest power of 1050 mW (Fig. 2b), 6 points of interest were selected and their fluorescence signal plotted over time ((Fig. 2c). None of the regions exhibited any detectable decrease in fluorescence over the recording period, indicating that — contrary to what might be expected based solely on excitation power — the 4D-SLIDE imaging regime does not induce observable photobleaching on this timescale. To check for signs of photodamage, we chemically fixed the tissue used for irradiation experiments with paraformaldehyde and inspected it by wide field fluorescence microscopy (Fig. 2a). A large area was investigated (several mm^2^), so as to include all SLIDE irradiated and several non-irradiated zones. The regions that were exposed to the irradiation are marked with white rectangles. We did not find any indication of damage or photobleaching (Fig. 2a), when adjacent SLIDE-imaged and non-irradiated ganglia (grouped neurons and glia) were compared. Throughout the tissue, the tdTomato fluorescence in the ENS appears homogeneous, with only naturally occurring variance in expression levels. This absence of photobleaching or photodamage appears remarkable at the powers employed, however similar trends were reported for rapid scanning systems before, possibly explained by a lower probability of transition to a triplet state upon picosecond excitation^13–15^. Further, the lower peak powers used with picosecond excitation pulses help avoiding nonlinear damage pathways^13,14^. As to the thermal load on the sample, we hypothesize that the fast scanning aids with thermal dissipation^16^. Based on these and subsequent measurements, we conclude that photobleaching or photodamage was absent or at least negligible as spontaneous tissue contractility was not affected by the imaging. Corroborated by these initial verifications, follow-up experiments in which the intestine was left tubular and intact were set up to acquire 3D image stacks of the multilayered ENS during spontaneous tissue movement (Fig. 3a).

**TABLE I:**
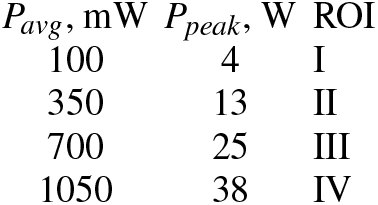
Average and corresponding peak SLIDE excitation power levels at the sample plane. In third column, the levels used in this study are associated to ROIs used in Results section (Fig. 2a).

**FIG. 2:**
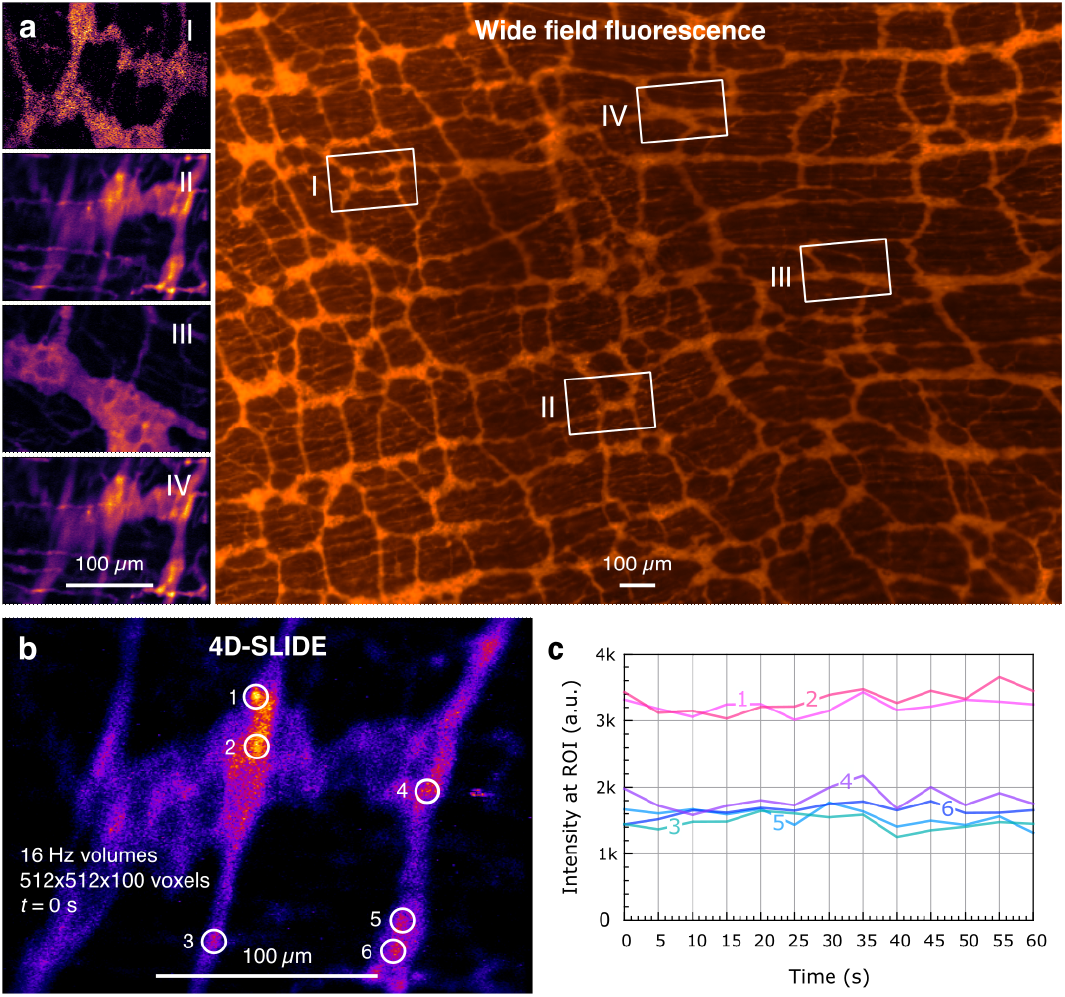
Light-tissue interaction measurement. **a** Four regions were imaged for 60s each at different power levels. The tissue was then inspected by widefield fluorescence microscopy. No Photodamage was found. For the highest power level used (*P*_*avg*_ = 1050 mW, see table I), the temporal evolution of the fluorescence signal was plotted **c**.

**FIG. 3:**
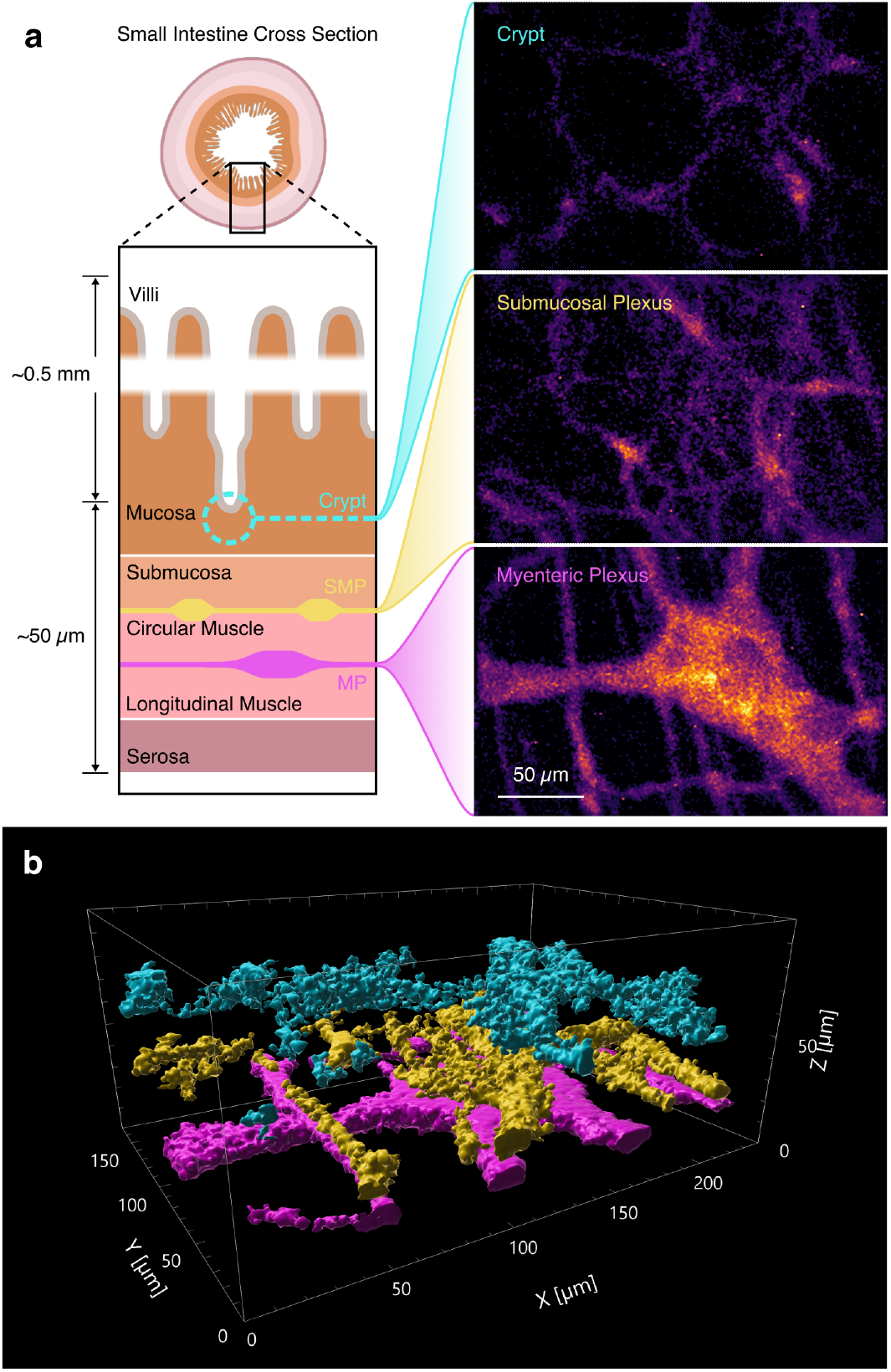
**a** Localization of the neuronal layers in the small intestine (left). Maximum intensity projection of the planes that include the cells at the bottom of the crypt, the submucosal plexus, and the myenteric plexus (right) acquired from a single 3D-SLIDE volume recording. **b** Volume rendered reconstruction generated using IMARIS software of the dataset in a. The myenteric plexus, the submucous plexus and the cell layer at the bottom of the crypt are color-coded in magenta, yellow and cyan, respectively.

### 4D-SLIDE enables simultaneous live 16 Hz volumetric imaging of the distinct neuro-glial layers and crypt-residing glia

Based on the characterization above, we selected *P*_*avg*_ = 700 mW for the imaging experiments in an excised, whole intestine tube. Imaging at this power level already provided sufficient signal intensity (see Fig. 2a. III). In a next set of experiments, we assessed the actual penetration depth of the 4D-SLIDE system. We used a fresh gut tube, not opened or peeled, from the same transgenic tdTomato expressing mouse. The fresh small intestine was placed in a holder with a glass coverslip bottom that was filled with oxygenated KREBS solution (see materials and methods). The intestinal tract has a concentrically layered structure composed of a serosal cell layer with connective tissue on the outside, and muscular, neuronal, and mucosal layers towards the lumen (Fig. 3a). Therefore, for imaging inside the intact intestine, the excitation beam needs to travel first through the serosa and the longitudinal muscle layer before reaching the myenteric plexus, the first fluorescent layer, which was easily detectable with 4D-SLIDE imaging (see Fig. 3). About 10-20 µm deeper and through the circular muscle layer, light reaches the submucosal plexus. Another 20 µm deeper inside the intestinal wall, we also recorded 2P-induced fluorescence from cells located at the bottom of the crypts (Fig. 3a, right). The real time rendering of the acquired volumes, capturing each of these three layers simultaneously, can be seen in Video 1 (left) accompanying Figure 3.

Next, we rendered the data obtained from 4D-SLIDE (Fig. 3a) in a commercial image analysis package (IMARIS) for further analysis. The volume rendering function easily identified each of the three surfaces, corresponding, respectively, to the myenteric plexus, submucous plexus, and the glial cells at the bottom of the crypts (Fig 3a). This showed that the signal fidelity is sufficient for segmentation, even at these high-speed volume rates. For easier visualization, we color-coded the layers by their position in z, in a short recording where the movement in z was minimal (Fig. 3b and Video 2).

Taken together, with the 4D-SLIDE system we simultaneously recorded 2P-excited fluorescence from relatively large volumes 250 × 150 × 100µm^3^, and thus a 310 µm 3D distance along the diagonal) comprising MP, SMP and glial cells present at bottom of the crypt. Here, the movement of the neuronal layers can be used as a proxy to describe the circular and longitudinal muscle layer contractions at the microscopic level, not only in 2D, but for the first time also in 3D. The ability to simultaneously resolve and follow the different layers in 3D holds promise for studying the control of these coordinated micro-movements, which are essential for mixing the luminal contents and facilitating absorption of nutrients^17,18^

### Live 4D-SLIDE microscopy allows extracting 3D spatio-temporal information about relative cell position and speed in intact and living intestine

In the next analysis, we set out to describe spontaneous gut movement, as observable within a 240 × 170 × 100µm^3^ volume. A sequence was selected (Video 3) of which three non-consecutive frames, are shown in Fig. 4a, with a tracked point speed indicated in the bottom left corner. One cell was identified and tracked over 90 frames. The trajectory of the tracked cell shows one cycle of the expected periodic movement (Fig. 4b), as known from standard wide-field imaging, but this time with resolved 3D components. The three-dimensional speed was calculated from the displacement over unit time (Fig. 4c). In Video 3 the colour coded line for the trajectory and the speedometer show the speed as a rolling average with a window size of *N* = 2. This was done to reduce the visual impact of Z-axis hysteresis, which caused alternating frames to exhibit systematically larger or smaller Z displacements during upward and downward scans, respectively. The 16 Hz volumes recorded with the 4D-SLIDE microscope provide good temporal and spatial resolution to describe the 3D trajectory of cells from the ganglia in the contracting gut tube. During the 6 s that the tracked cell stayed in the imaged volume, its maximum displacement was 221 µm in x, 121 µm in y and 25 µm in the z direction and its speed ranged from 30 to about 240 µms^−1^ (Fig. 4c).

**FIG. 4:**
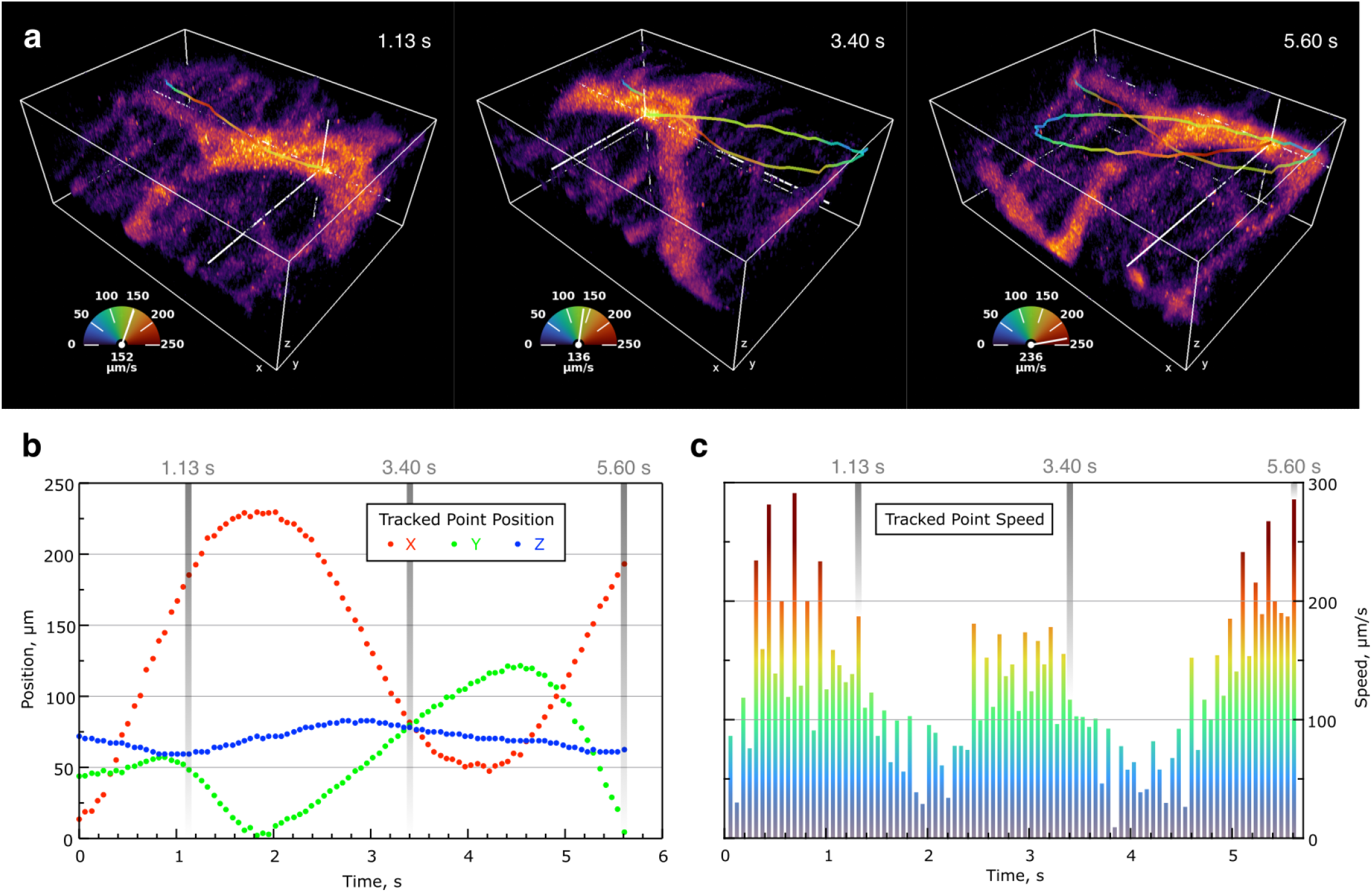
Tracking cells in the contracting gut. **a** Individual snaphsots of a 4D-SLIDE dataset. The position of the feature indicated by the white crosshair was manually tracked over time. The colors in the trajectory trace correspond to the instantaneous speed according to the speed gauge in the lower left corner. **b** *x, y, z* position (left) and calculated speed (right) of the tracked feature.

### 4D-SLIDE enables measuring relative 3D movement between the two muscular layers of the intestine

Being able to monitor neuro-glial structures in the intact gut, at sufficient speed and with the optical sectioning capacity of 2P excitation to resolve the individual layers, we investigated if 4D-SLIDE could also resolve micrometer differences in how the different layers of the intestine move in 3D. This would make it possible, for the first time, to study distortions of neuro-glial layers as the intestine contracts in normal physiological operation.

We identified a distinguishable spot in the myenteric and another one in the submucous plexus and tracked their movement over time (Fig. 5a-c magenta and yellow spheres, respectively). As in the previous recordings, the direction of the longitudinal muscle layer is quasi parallel to the direction of the X axis, while the circular muscle direction is aligned with the Y axis. Selected frames from a 4D-SLIDE acquisition (Video 4) are presented in Fig. 5d. Here, the contractile tissue is seen moving in 3D across the imaged volume. The trajectories of the two tracked spots in the myenteric and submucosal plexus layers (Fig. 5d) are shown as traces.

**FIG. 5:**
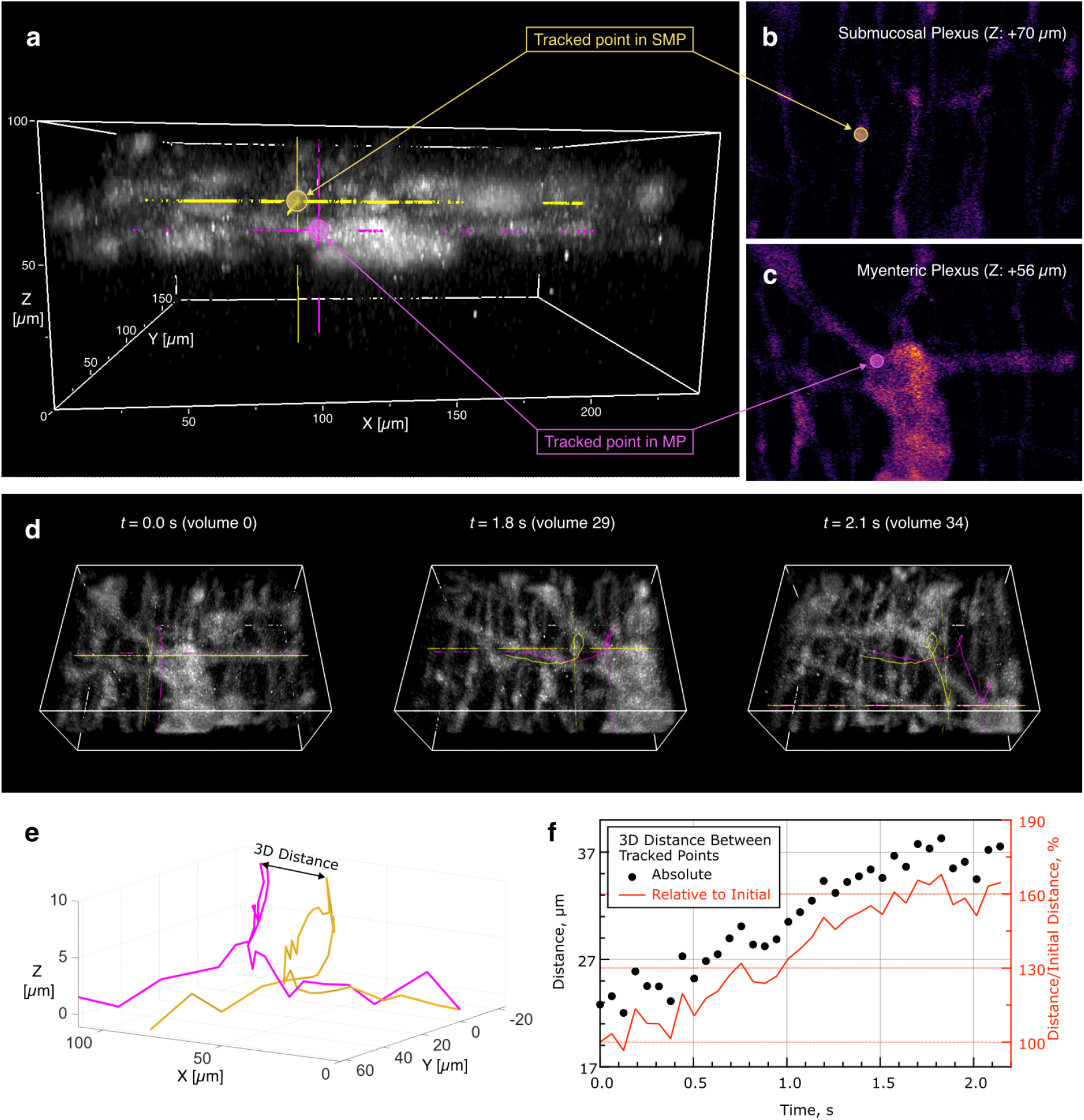
Micrometric independent displacement of the intestinal muscular layers. **a** Side view of MP and the SMP with the tracked object indicated in each. Optical section of the SMP (**b**) and MP (**c**). The yellow and magenta dots indicate the tracked features in the two layers. **d** The magenta and yellow traces show the trajectories of the spots in b and c. **e** Dimetric plot of the trajectories over time highlighting the 3D distance at a given time. **f** Time-evolution of the absolute (black) and relative to the initial position (red) 3D distance.

In the initial frame, the 3D distance between the tracked spots is 23 µm, and should not change significantly if both layers would move together. However, the 3D distance grows, reaching a max of 38 µm at frame 30 (Fig. 5f, *t =* 1.87 s). Such 15 µm difference represents a 65 percent change over the initial distance. The dimetric 3D plot of the two trajectories (Fig. 5e) shows a similar but not identical movement pattern for the two layers, mostly pronounced laterally (longitudinal muscle contraction or relaxation), as the separation in axial direction stays largely unchanged for the two layers. In addition, both X and Y trajectories exhibit a non-uniform evolution: The tracked spots stop, move back, and resume transit across the scanned volume in initial direction. At their fastest speed, the tracked points move with a velocity of 150 µm/s in *X* and 260 µm/s in *Y* .

## III. DISCUSSION

In this paper, we presented the first physiological application of 4D-SLIDE microscopy for volumetric two-photon imaging. The 4D-SLIDE microscope relies on a diffraction based scanning approach to achieve unprecedented imaging speeds, generating 4D information much faster than other scanning techniques. Furthermore, the 2-photon excitation principle enables imaging deep inside opaque living tissue, where classic single photon (lightsheet, confocal) excitation would fail. We used mouse small intestine and imaged the distinct neuro-glia layers in the intestinal wall, taking advantage of a genetic expression system to selectively express the tdTomato reporter only and specifically in neurons and glia. The tubular sample, removed from the animal, was left intact to allow all natural contractions to occur in 3 dimensions. By consequence of their complexity (not transparent, non-uniform and contractile), imaging inside these tissues has not been possible without impactful interventions, either relying on physical reduction (stretching, dissecting) or pharmacological silencing of the contractile activity^19,20^. The ability to image inside the organ, while leaving it intact and freely contracting, opens up important research avenues for studying the effects of distortions and local stretching of muscle, neuro-glial connections and extrinsic innervation.

The excitation source in 4D-SLIDE is, by design, operated at duty cycle orders of magnitude higher than those used in fs-2PM - both because of the longer pulse duration (30 ps), and because of the very high voxel rate (820 MHz). A detailed comparison of the excitation regimes used in SLIDE and in conventional fs-2PM has been presented in previous work^21^, where we defined the Pixel Excitation Integral (PEI) as a figure of merit for the number of photons generated from a single pixel:

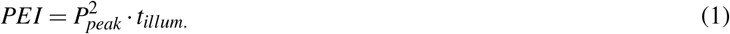

with *P*_*peak*_ as the peak power, and *t*_*illum*._ as the pixel illumination time. Here *PEI* = 0.019 kW^2^ ps for imaging in living tissue, which corresponds to *P*_*peak*_ = 25 W, and *P*_*avg*_ = 700 mW. In comparison, conventional fs-2P imaging lasers operate with average powers in the tens of mW^22,23^ and peak powers in excess of ∼1 kW^3^ and, therefore, PEI tens of times higher than in the case of SLIDE. The substantially lower PEI of SLIDE therefore necessitates the use of bright probes for fast imaging. While operating at a pixel dwell time of ~1 ns the total number of photons available from a voxel during a single scan is lower in SLIDE. The significantly lower *P*_*peak*_ suggests a reduced rate of photobleaching, as can be seen in Fig. 2c. At the same time, the average power in SLIDE is higher than in fs-2PM (see Table I), and a correspondingly increased thermal load can be expected. While this may be the case, it did not result in observable tissue damage or a physiological impairment. It is possible that in our experiment tissue integrity had a protective effect, aiding with heat dissipation, on the one hand but also the fast volumetric quality of 4D-SLIDE avoids cumulative focal heating in principle.

One of the current limitations of the technique relates to its intrinsic own advantage, that is its unprecedented scanning speed, leaving only very little time to collect photons from the excited spot. Therefore, at present only probes with a high quantum yield can be used (e.g. tdTomato), to ensure that a sufficient signal to noise ratio is reached. We show that, indeed, the generated 4D image stacks are of sufficient quality for direct and interactive interpretation on the screen, interoperable with existing analysis tools (e.g. tracking), as well as for rendering purposes in widespread commercial 3D visualization software.

One of the important advantages of the 4D-SLIDE imaging modality is that acquired data are rendered and displayed with latency lower than operator reaction time (<100 ms). As such, this allows to navigate with a 3D volume view in the sample while it is active (in this case contracting). This improvement in data collection and visualisation is in principle comparable, yet much more impactful, to the transition from slowly refreshing confocal image display to the ability to have live 2D image updates when focusing in tissue with spinning disk or resonant scanner confocal microscopes. The possibility to now have 3D volumetric information on the fly is transformative as it allows even better interpretation of the structures while imaging is ongoing and real-time interaction with living cells and tissues. In this ENS imaging example the volumetric acquisition frame rate was set to 16 Hz and the image size to 240 * 170 * 100 µm (512x1024x100 voxels), which enables the user not only to observe new features coming into the scanned volume as the tissue contracts, but also to react, toggling the recording of the acquired data or manipulating the stage to follow the object as it moves out of the scanned volume.

With the ability to simultaneously observe and record volumetric data at tens of volumes per second, a set of new research questions can be addressed. The importance of the biomechanical dynamics and the local forces that are exerted on the intestinal layers have yet to be explored. We were able to record with cellular resolution the two neuronal layers as they were moving due to circular- and longitudinal muscle contractions of the intact intestinal tube. Moreover the ability to resolve the two layers at this fast timescale, allows detecting that both layers do not move identically, which suggest they are experiencing different stretches. Already in the particular example presented in this paper, in which the compound movements of the tdTomato-labeled ENS serve as a proxy for the contractions of the circular and the longitudinal muscle layer, we detect that both layers are skewed during normal physiological contractions. The impact that these deformations have on the local neurons and their connections can now be investigated in 3D and in the intact organ. Studies applying pressure stimuli in the enteric nervous system have shown that subpopulations of enteric neurons are responsive to local induced deformations^24^. For these experiments, intestinal preparations need to be dissected and pinned flat in order to be accessible by classic microscopy techniques. Even though these flattened preparations have yielded invaluable data, including information about the specific and independent innervation of the different muscle layers and their intracellular Ca^2+^ handling^25^, the ability to record contractions and distortions in 3D constitutes an important step forward. 4D-SLIDE microscopy now allows to start addressing these pressure- and stretch-related phenomena within the intact organ.

Apart from recording the mechanical motion at high spatio-temporal resolution, further investigation of how the ENS integrates information to produce functional output requires the ability to record neuronal activity. This is mostly accomplished using genetically-encoded calcium reporters, namely GCaMP. Unfortunately, these green-fluroescent proteins cannot be excited using the current 1064 nm laser. The development of a dedicated excitation source for green fluorescent proteins will allow full exploitation of SLIDE’s 4D imaging capabilities, enabling the recording of neuronal, glial, and muscle activity through the expression of GCaMP indicators in distinct cell types,^19,20,26,27^ while preserving intact, freely moving organ preparations.

In conclusion, we demonstrate the first application of 4D-SLIDE microscopy for live volumetric two-photon imaging inside intact, contractile tissue, without apparent photobleaching or photodamage. By leveraging akinetic scanning to achieve MHz line scan rate, 4D-SLIDE combines video-rate volumetric acquisition and deep two-photon penetration for biological tissue imaging. Such an operating regime is inaccessible to conventional femtosecond two-photon microscopy due to volumetric scan speed and pulse repetition rate limitations. While current SLIDE excitation source favours bright red fluorescent probes, ongoing developments in excitation wavelength, detection efficiency, and fluorophore compatibility are expected to further broaden applicability. More broadly, 4D-SLIDE introduces an architecture for high-speed, deep, and interactive volumetric imaging, with potential impact across biological, biomedical, and dynamic material systems.

## IV. MATERIALS AND METHODS

### A. 4D-SLIDE microscope

The 4D-SLIDE microscope is built on the platform of an inverted fluorescence microscope (*Nikon Ti2 Eclipse*). SLIDE enables kHz frame rates and MHz line rates by employing a wavelength-swept, pulsed excitation source^11^ in combination with a diffraction grating in the fast axis (akinetic spectro-temporal scanning). A Fourier-domain mode-locked (FDML) laser^28^ operating at 1064 nm central wavelength (*Optores NG-FDML*) is used as the swept source. The continuous-wave emission of the FDML laser is modulated using a high-bandwidth electro-optic modulator (EOM), forming a *λ* -swept train of 30 ps long pulses, with a 820MHz pulse repetition rate. The pulses are then amplified in a two-stage ytterbium-doped fiber amplifier, providing the peak power necessary for two-photon excitation (see table I). The sweep is pre-modulated to compensate for ytterbium gain variation^29^ and provide a consistent output power level for even illumination. The excitation pulse power is controlled by adjusting the gain of the final stage of the ytterbium-doped fiber amplifier via the pump laser power. The average power (*P*_*avg*_) was measured (*Thorlabs PM100D, S170C*) at the sample plane during on-site experiments. The pulse peak power (*P*_*peak*_) is related to the average power in Table I. The scanning along the *Y* axis was performed using a galvo scanner (*Scanlabs dynaxis421*), and the focal plane is scanned along the *Z* axis using a piezo objective scanner (*Thorlabs PFM450E*). We chose ∼ 16 Hz volumetric acquisition rate (63 ms per acquisition) as optimal to match the axial scan range (100 µm) with the desired sampling density.

#### 1. Mapping, sampling density and PSF

Using a 25x NA1.1 objective (*Nikon N25X-APO-MP1300*) we scanned a field-of-view (FOV) of 240 × 170 × 100 µm (*XYZ*) in size, which was sampled in 512 × 1024 × 100 voxels (see table II). To correct for non-linearities of scan (sinusoidal scanning patterns), the volumetric data is later resampled in post-processing to a final resolution of 461 × 652 × 64 voxels of uniform size (see detailed sampling information in Table below). The sampling factor (point spread function divided by voxel size) along *X* and *Z* was selected to be lower than 2 to achieve desired scan speed. Additionally, the fluorescence lifetime of TdTomato can be 2.9 ns to 3.5 ns^30^, which already exceeds the 1.2 ns long voxel dwell time, and the signal of decaying fluorescence is already carried over into the adjacently scanned voxels. A larger FOV at the expense of sampling density in both *X* and *Z* was deemed beneficial for the actual experiment to ensure that a sufficient number of structures (ganglion, connecting fiber bundles) stayed within the scanned volume during contractile motions of the living tissue.

**TABLE II:** Scan field and Sampling Table.

| Axis | X | Y | Z |
| --- | --- | --- | --- |
| FOV size, $\mu\text{m}$ | 240 | 170 | 100 |
| Voxel count | 512 | 1024 | 100 |
| Voxels after numerical linearization | 461 | 652 | 64 |
| PSF (measured), $\mu\text{m}$ | 0.72 | 0.64 | 1.28 |
| Voxel size (calculated), $\mu\text{m}$ | 0.52 | 0.26 | 1.56 |
| Oversampling factor (2 is Nyquist limit) | 1.38 | 2.46 | 0.78 |

#### 2. Signal Acquisition and Processing

The two-photon fluorescence signal is acquired using a hybrid photo-detector (*Hamamatsu R11322U-40-01*) and amplified by a high speed trans-impedance amplifier (*Femto HCA-400M-5K-C*). The signal is then processed by a high-speed 12 bit digitizer card (*AlazarTech ATS9373*), where it is sampled at 3.3 GSa/s (sampling rate phase-locked to 4 samples per voxel or 2048 samples per FDML sweep). The resulting data stream, reaching up to 5 GB/s, is handled by custom software for real-time 3D rendering of the volumetric 2PM data, with the option to simultaneously record it (Fig. 1c). The recorded raw data is compressed and stored in a custom format. *Tiff* files are generated from the raw data to facilitate compatibility with common image processing software (Napari, Fiji, Imaris, etc.). The recorded volumes are also filtered using a Python implementation of the 3D hybrid median filter^31^ to reduce the background and spurious noise. A comparison of filtered and unfiltered data is presented in Video 1. Imaris software (*Bitplane, Oxford Instruments*) was used to generate the surfaces shown in figure 3 and video 2. A short recording where the movement of the intestine was mainly along the *X* and *Y* directions was selected to generate surfaces based on a fixed range in the *Z* position of each intestinal layer. In figures 4 and 5, different features in the sample are followed over time. Since tdTomato is highly expressed in many cells, which are optically overlapping, automated tracking algorithms produce inaccurate results. Therefore, the position of the selected cell was manually determined in Imaris at each time point. The position data of the different features is then used to calculate the motion speed (figure 4) or the distance (figure 5) of the tracked features. The Data is visualized in Napari, using an adjusted ‘inferno’ LUT with a gamma value of 0.5. Traces are added to visualize the trajectory of the tracked spots.

### B. Tissue preparation

Heterozygous Wnt1-Cre mice were crossed with R26-lsl-tdTomato reporter mice, resulting in offspring (in short: Wnt1|tdTomato), in which the neurons and glia cells of the peripheral and enteric nervous system were expressing the red fluorescent protein tdTomato. These Wnt1|tdTomato mice were kept in filter-top cages in a controlled environment with room temperature between 20 °C to 21 °C, humidity between 50 % to 60 %, and a light–dark cycle of 12 h. Just before the imaging experiments, the animals were killed by cervical dislocation, after death was confirmed, the intestine was removed from the abdomen and kept in oxygenated buffer (see below). All procedures were conducted in accordance with the ethical standards for experiments on animals established and approved by the Animal Ethics Committee of KU Leuven.

Two types of tissue preparations, were used in this study: on the one hand a stretched gut tissue preparation and on the other hand a whole intestinal tube, which were both kept alive in oxygenated (5% O_2_ to 5% CO_2_) Kreb’s solution (containing in mM: 120.9 NaCl, 5.9 KCl, 1.2 MgCl_2_, 1.2 NaH_2_PO_4_, 14.4 NaHCO_3_, 11.5 glucose, 2.5 CaCl_2_) at room temperature.

The stretched tissue was important to test the effect of average power and therefore the distal ileum was dissected and pinned flat in a Sylgard-lined dish filled with Kreb’s buffer. The mucosa and submucosal layers were carefully removed using microdissection, to generate a preparation comprising longitudinal, circular muscle and the embedded myenteric plexus. This tissues was mounted over a small inox ring (5 mm inner diameter) and immobilized by a matched rubber o-ring^9,32^) and further pharmacologically inhibited by nifedipine (2 µM), as to maximally restrict the endogenous contractile movement and enable irradiation of exactly the same region. In a second preparation, the intestinal tube (proximal ileum) was kept intact, and gently flushed with Kreb’s solution to remove the luminal contents. A segment of intestine, approximately, 1.5 cm long was mounted over an inox ring (1 cm inner diameter) and immobilized by a matched rubber O-ring. The latter preparation, with all its cellular layers, architecture, morphology and contractility intact, is an ideal physiological sample to assess the 3D imaging capabilities of the fast 2P imaging system. Indeed, the intestine with its embedded enteric nervous system (ENS) remains alive and muscle (circular and longitudinal) contractions are still ongoing, continuously distorting the tdTomato-labeled ENS. Recordings were made at room temperature (22 °C).

### C. Photobleaching and Photodamage

To analyze photobleaching in 4D SLIDE acquisition, we recorded a 60 s continuous illumination (16Hz volume rate) on the stretched tissue preparations at maximum power level (*P*_*avg*_ = 1050 mW) . From the dataset we extracted a sample volume every 5 s and z-projected (maximum intensity projection) into 2D intensity maps. Small ROIs were drawn and manually adjusted for each time point to match selected features as they move within the FOV over the duration of acquisition (Fig. 2b). The signal intensity of each ROI was then plotted over time (Fig. 2c), showing no apparent reduction over the duration of 1 minute. To analyze visually the imaged areas, the tissues were paraformaldehyde fixed and then imaged using a widefield microscope (*Zeiss Axio imager*). The entire stretched tissue area was imaged and then the regions exposed to SLIDE irradiation were located in the fixed tissue (cf. Fig. 2a). We did not observe bleaching of the irradiated areas by widefield imaging.

## Supporting information

No extra supplementary information

## Acknowledgements

S.K., LB and PVB acknowledges funding from EU Horizon2020: FAIR CHARM, grant 101016457. SK further acknowledges support from the Deutsche Forschungsgemeinschaft (EXC 2167-390884018, 511288691, project ADAPT - KA 4354/6-1, KA 4354/7-1) and EU research funding project SWEEPICS, grant 101135053. LB acknowledges funding from SNF (200021E-213168 MINT) for ADAPT, and PVB acknowledges FWO support for ADAPT (G070223N) as well as KU Leuven Methusalem (METH/014/05) support.

